# Host selection has stronger impact on leaf microbiome assembly compared to land-management practices

**DOI:** 10.1101/2023.03.02.530479

**Authors:** Pankaj K. Singh, Eleonora Egidi, Catriona A. Macdonald, Brajesh K. Singh

## Abstract

Plant microbiome contribute directly to plant health and productivity but mechanisms that underpin plant microbiome assembly in different compartments (e.g. root, leaf) are not fully understood. Identifying environmental and management factors that affect plant microbiome assembly is important to advance understanding of fundamental ecological processes and harnessing microbiome for improved primary productivity and environmental sustainability. Irrigation and fertilization are two common management practices in Australian tree plantations, but little is known about the effects of these treatments on soil, plant host, and their microbiome. Here, we investigated the impact of a decade long irrigation, fertilization, and their combined application, on soil, plant traits and microbiome of a *Eucalyptus saligna* plantation at the Hawkesbury Forest Experiment, Western Sydney University, Richmond, NSW. Microbial profiling of bulk soil, rhizosphere, root, and leaves was performed using amplicon sequencing 16S rDNA and ITS markers for bacteria and fungi, respectively, along with measurements of soil properties and plant traits. The results indicated that both management practices affected soil properties and soil and root microbiome significantly. Irrigation increased but fertilizer treatment reduced microbial alpha diversity. However, neither irrigation nor fertilizer treatment significantly impacted the leaf microbiome. Our findings imply that management practices impact soil edaphic factors, which in turn influence the below ground microbiome (soil and root). In addition, the leaf microbiome was distinct from soil and root microbiome, and a source tracker analysis suggested root and bulk soils only contributed to 53% and 10% OTUs of the leaf bacterial community, suggesting strong and sequential host selection of the leaf microbiome. In addition, management practices had limited impact on leaf traits and, consequently, the leaf microbiome maintained its distinct composition. These findings provide mechanistic evidence for ecological processes that drive plant microbiome assembly and indicate that host selection plays a more important role than management practices on leaf microbiome assembly.

## Introduction

The plant microbiome is considered the extended part of a plant’s genome and plays a pivotal role in regulating plant fitness and functions. For example, the microbiome provides resilience against biotic and abiotic stresses in exchange of energy and habitats provided by plant hosts (Trivedi *et al*., 2020). This association is mutually beneficial which has coevolved over millennia(Heckman *et al*., 2001). Current paradigm suggests that the microbiome assembly in plants is regulated by a plethora of biotic and abiotic factors, ranging from soil-associated variables that impact soil microbiome such as soil pH, moisture, and nutrient availability (Turner *et al*., 2013; Compant *et al*., 2019), to host-associated factors such as host genotype, physiology, and plant compartment (Hamonts *et al*., 2018; Xiong *et al* 2021). It is proposed that the host plant employs a filtering mechanism to select microbial endophytes (Xiong *et al*.,2020). In general, the host plant releases chemicals and root exudates into the soil which serve as a signal to initiate the recruitment of specific microbes in the rhizosphere, which subsequently enter inside the plant through the root and are then transported across the plant compartments through shoot into leaf, seeds, flower, etc. (Miransari and Smith,2014; Shade *et al*.,2017, Liu *et al*., 2020). The plant microbiome assembly requires mutual recognition where microbes recognize chemical signals produced by the plant to respond, while the plant immune system ensures that only recognized microbes can colonize (Fitzpatrick *et al*., 2020). However, relative contributions of different ecological processes (e.g. roles of host vs environmental vs managements factors) that govern plant microbiome assembly remain poorly understood.

Other environmental factors such as management practices, the presence of pathogens, and climate are also known to play roles in the plant microbiome assembly (Liu *et al*., 2017, Ware *et al*., 2021). For example, land management practices such as irrigation, tillage, use of fertilizer, and pesticide, can influence plant growth and survival. These management practices are also known to affect soil and rhizosphere microbiome assembly (Kavamura *et al*., 2018; Hartman *et al*., 2018; Toju *et al*., 2018; Sengupta *et al*., 2020). For example, nitrogen fertilizers were reported to reduce bacterial diversity (Zhang *et al*., 2018; Yang *et al*., 2020). On the other hand, fertilizer application has been observed to improve both above and belowground plant fungal diversity (Fornara and Tilman, 2012; Li *et al*., 2015). Irrigation, on the other hand, increases the soil moisture content and soil microbial diversity (Colombo *et al*, 2016; Zheng *et al*., 2017). Recent studies have demonstrated increased bacterial and fungal diversity under irrigated sites but not under fertilization treatments (Colombo *et al*, 2016;

Zheng *et al*., 2017) which was linked to increased rate of soil functions (Colombo et al. 2016). However, the response of plant (e.g. root, leaf) microbiome to land management practices is not known. This is a major knowledge gap given the soil microbiome is considered as the main source of plant microbiome which ultimately affects plant health and productivity (Singh *et al*., 2020; Trivedi *et al*., 2022). Such knowledge is needed to develop microbial solutions to increase plant productivity in environmental sustainable way (Li *et al*., 2022; Trivedi *et al*., 2022)

In this work, we aimed to examine the impact of management practices (irrigation and fertilizer) on *Eucalyptus saligna* (commonly known as Sydney Blue Gum and widely used by the plantation industry) microbiome assembly. We hypothesized that (i) management practices will affect the soil edaphic properties and soil microbiome assembly, in turn, it can influence the diversity and structure of endophytic (e.g root and leaf) communities, (ii) leaf endophytes will be less impacted by management practices compared to root microbiome due to host specificity and sequential host filtering. To test these hypotheses, we analyzed the bacterial and fungal composition of four different niches, bulk soil, rhizosphere soil, root, and leaves of *Eucalyptus saligna* under irrigation, fertilizer, and irrigation plus fertilizer treatments. Further, we measured leaf traits to account for management induced changes on traits and its impact on leaf microbiome. To understand the implication of host-associated factors and their effect on plant microbiome assembly, we built a plant microbiome source prediction model and analyzed the impact of soil and leaf-associated factors on the community assembly..

## Material and methods

### Sampling site and sampling strategy

The field site is located at the Hawkesbury Forest Experiment (33°36’40’’ S, 150°44’26.5’’ E), at Western Sydney University, Hawkesbury Campus, Richmond New South Wales. It was converted from natural to improved pastures in 1997. The site is comprised of an alluvial formation of sandy loam soil classified as a Chromosol, with low soil organic matter content (0.7%), and low water holding capacity (Barton *et al*., 2010; Hu *et al*., 2015). *Eucalyptus saligna* were planted in 2007 at a density 1000 trees per hectare. The site has a total of 16 plantation plots. The mean annual temperature of the site is 16.8 ° C and annual precipitation is about 870mm per year (https://en.climate-data.org/oceania/australia/new-south-wales/richmond-14607/).

The experiment consisted of four different treatments; an unaltered condition i.e., without application of irrigation and fertilizers (Control-C); a fertilizer treatment (F), an irrigation treatment (I); a fertilizer x irrigation (IF). Sixteen plots were arranged in a randomized block design with four blocks each containing four treatment plots, each measuring 38.5 × 41.6 m. Trees were planted in 10 rows for each plot. At the time of planting, each plant was treated with an application of 50 g of diammonium phosphate blend (nitrogen [N] 15.3%, phosphorus [P] 8.0%, potassium [K] 16.0%, sulphur [S] 7.7%, and calcium [Ca]0.3%) to promote growth. Control plots received no further irrigation or fertilizer application. For both F and IF treatments, the first application of solid N fertilizer took place in January 2008 (N 20.6%, P 3.0%, K 7.5%, S 3.8%, and Ca 4.4%) at an amount equivalent to 25 Kg N ha^-1^year^-1^. From October 2008 onwards, solid N fertilizer (N 21.6%, P 8.1%, K 12.0%, and S 0.6%) application was initiated for F treatment at a rate of 150 Kg N ha^-1^year^-1,^ and IF plots were treated with liquid fertilizer 150 Kg N ha^-1^year^-1^ (Nutrifeed19 and Liquid N, Amgrow Fertilisers, Lidcombe, NSW, Australia). Both I and IF treatments received water at a rate of 1000 mm year^-1^ in addition to rainfall (Zheng *et al*., 2017).

Sampling was conducted twice to check for temporal variations, the first in the Autumn (February 2020) and repeated in the following spring (October 2021). In 2020, three healthy trees from each of the 16 plots were selected from the middle of the plot to avoid any edge effect. Around each tree, three soil cores (2.5 cm wide and 15 cm deep) were taken. The replicate soil samples for each tree were pooled and *E. saligna* roots were separated from the soil. Soil samples were sub sampled. For soil physicochemical properties, samples were sieved through 2mm mesh and stored at 4°C, while other soil and root samples were stored at -20°C for DNA extraction.

Using a cherry picker, young green and healthy leaves from each of the selected trees were collected for leaf trait and nutrient analysis as well as for DNA extraction from a uniform height of 15 m. Leaves for DNA isolation were stored at -20°C.

### Soil Physico-chemical properties

Soil pH was determined by shaking 2.5±0.01 g of soil with 12.5 ml of MilliQ water at 180 rpm (Lab Companion SK-71 Shaker, USA) for 1 hour at room temperature after which pH measured using a Delta pH meter (Mettler-Toledo Instruments, Columbus, OH, USA). Soil moisture was determined by mass loss after drying at 105° C for 24 hours. Soil moisture content was determined as a percentage of water per gram of dry weight of soil.

Extractable inorganic N was determined on fresh bulk soil by shaking soil with 2M KCl (1:10 w:v) for 1 hour at 180 rpm. Extracts were filtered (Whatman 42) and analyzed colorimetrically for determination of nitrate (NO_3_) and (NH_4_) (Keeney and Nelson, 1982) using a SEAL AQ2 Analyzer (SEAL Analytical, Maquon, WI, USA). Extractable inorganic P was determined on fresh bulk soil by shaking with Bray’s reagent (0.03 M Ammonium fluoride - NH_4_F and 0.025 M Hydrochloric acid -HCl) (Bray and Kurtz, 1945) for 1 minute at 180 rpm. Extracts were filtered (Whatman 42) and analyzed colorimetrically using a SEAL AQ2 Analyzer (SEAL Analytical, Maquon, WI, USA). Over-dried soil samples were ground using a Mixer Mill MM400 grinder (Retsch, Germany), prior to determination of total C and N using a LECO macro-CN analyzer (LECO, St Joseph, MI, USA).

### Measurement of Leaf functional and chemical traits

Leaf dry weight was determined following drying at 40°C for 48 hours. Leaf dry matter content (LDMC, mg. g^-1^.) determined using the following formula:

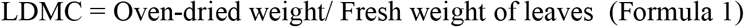

Specific leaf area (SLA; mm^2^.mg^-1^) was determined using 5 dried leaves for each tree and was calculated using the following formula (Pérez-Harguindeguy *et al*., 2016) :

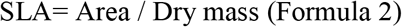

Total C and N were obtained from oven-dried leaf samples as method as described for total C and N for soil.

### Surface sterilization, DNA isolation, and amplicon sequencing

Rhizosphere soil was collected by washing roots with sterile Milli-Q water. Root and leaf samples were surface sterilized with 70% ethanol for 2 minutes, followed by 2% sodium hypochlorite for 3 minutes, and again with 70 % ethanol for 1 minute. The samples were washed twice with sterile distilled water to remove any remnants of sterilizing agents. After surface sterilization, the leaf and root samples were cut into small pieces. DNA extraction was performed on 0.25 gm of leaf, root, bulk soil, and rhizosphere soil samples using the PowerSoil^®^ DNA Isolation kit (Qiagen Inc., Valencia, CA, USA), following the manufacturer’s instructions. The quality of DNA was evaluated using a NanoDrop spectrophotometer (NanoDrop Technologies, Wilmington, DE, USA). Microbial community profiling was performed using amplicon-based sequencing methods. For bacteria, the V5-V6 region of the 16S rRNA gene was amplified using 799F and 1193R (Kembel SW *et al*., 2014) set of primers, whereas for fungi, Internal Transcribed Spacer 2 (ITS2) region was amplified using fITS7 and ITS4 primers (Ihrmark K, *et al*., 2012) at the Next Generation Sequencing Facility at Western Sydney University (Richmond, New South Wales, Australia) using Illumina MiSeq sequencing platform.

### Bioinformatic Analysis and data processing

Sequenced raw reads were initially processed using FASTQC (Andrews 2010) for quality control and high-quality reads having a Quality score(Q) > 30 were processed using Mothur V1.46.0 (Schloss *et al*., 2009, 2020). Unique sequences were selected for processing and chimeric reads were identified using VSEARCH and eliminated from the analysis. Non-chimeric reads were then classified using Mothur’s Bayesian classifier for taxonomic classification. Unite (V8.2) and Silva (V138.1) databases were used for fungi and bacteria, respectively. The sequences were picked at 97% similarity. All mitochondrial and chloroplast-associated classifiers were removed, and the remaining reads were classified into Operational Taxonomic Units (OTU). All singletons were removed from the analysis following the OTU assignment to get rid of errors due to sequencing artifacts. Altogether, 3.4 million reads for bacteria and 8.5 million reads for fungi were obtained. The processed files were then used for statistical analysis. Samples were rarefied at 400 reads for analysis (for leaf) as a minimum library size for both fungi and bacteria. Samples were then rarefied for individual plant compartments, 1,000 reads per sample for root and samples were again subset at 10,000 reads for both bulk and rhizosphere soil. Though, no significant differences were observed when samples were rarefied for individual compartments. (Supplementary figure 1,2,3 and 4)

### Statistical Analysis

Files generated from Mothur were imported into R and processed using the phyloseq package (McMurdie and Holmes, 2013). Samples were subset niche-wise (bulk soil, rhizosphere, root and leaf) and analysed. The alpha diversity index (Shannon index) was calculated using the VEGAN package (Okansen *et al*., 2020), and differences in alpha diversity across treatments were visualized as box plots using ggplot2 (Wickham H., 2016). A negative binomial generalized linear mix model (nbGLMM) analysis was performed to identify major drivers of bacterial and fungal alpha diversity using the R package (R Core Team, 2021) MASS (Venables and Ripley, 2002) for each niche. The Beta diversity of both bacterial and fungal communities was assessed using the Bray-Curtis distance matrix and further ordinated using Principal Co-ordinate Analysis (PCoA) using Vegan and ggplot2 packages. The relative contributions of variables such as treatment, time point, soil, and leaf-associated factors were evaluated with PERMANOVA for every niche at 999 permutations using a Vegan package. Additionally, Spearman correlation analysis was performed to explore the relationship and impact of management practices, soil and leaf-associated traits and factors, and microbial diversity. The result was represented in form of a correlogram. One-way ANOVA and Tukey’s post hoc test were performed to assess the effects of management practices over soil measures and leaf traits. For ANOVA, management practices were used as predictor.

Differential abundance analysis was performed using the edgeR package (Robinson *et al*., 2010). Samples were first divided into subsets according to plant niches- leaf, root, rhizosphere soil, and bulk soil. Then, OTUs prevalent in more than 50% of samples were selected for further analysis. Subsequently, the samples were analyzed across all treatments, and only significantly enriched OTUs (P-value >0.05) were retained(Nearing *et al*., 2022).

The Source Tracker (Knight *et al*, 2011) tool from QIIME(V.1.9.0)(Caporaso *et al*., 2010) was used to build a source prediction model. The OTU table and associated metadata files were imported into QIIME for this analysis. This tool uses the source and sinks approach to detect the possible source of OTUs based on metadata description. The following niches were selected as a source at different instances

Using the information from source tracker analysis, plant microbiome source prediction models were designed for both fungi and bacteria.

## Results

### General description of the dataset

We found that the phylum-level composition of the plant microbiome differed substantially between different plant niches (Supplementary figure 5 and 6). The bacterial community was represented by a total of 22,784 OTUs, divided among the leaf (3,628 OTUs), the root (8,021 OTUs), the rhizosphere soil (13,625 OTUs), and the bulk soil (16,384 OTUs). The bulk soil was dominated by P*roteobacteria* (39%) followed by *Actinobacteria* (24%), *Firmicutes* (13%), and *Acidobacteria* (6%). In the rhizosphere, *Actinobacteria* accounted for nearly 35% of the bacterial community followed closely by *Proteobacteria* (31%), *Firmicutes* (8%), and *Acidobacteria* (5%). This trend was different in root endophytes where *Proteobacteria* accounted for 45% of the bacterial community followed by *Actinobacteria* (38%) and *Acidobacteria* (4%). *Proteobacteria* was more abundant in leaf compared to other Phyla, accounting for >62% of the bacterial community followed by *Actinobacteria* (21%) and *Firmicutes* (8%).

The fungal microbiome, or mycobiome, had a total of 18,717 OTUs. Compartment-wise leaf had 2,397 OTUS, the root had 4,163, the rhizosphere had 9,947 OTUs and the bulk soil had 12,487 OTUs. Contrary to the bacterial community, the phylum-level composition did not change substantially in terms of dominance across the compartments (Supplementary figure 6). The leaf mycobiome was dominated by *Ascomycota* (80%), *Zygomycota* (7%), and Basidiomycota (2%). Root had a similar trend in terms of dominance, but the composition varied here, *Ascomycota* (71%), *Zygomycota* (22%), and Basidiomycota (2%). In the rhizosphere, *Ascomycota* accounted for nearly 78% of the fungal population, followed by *Zygomycota* (15%) and Basidiomycota (2%) whereas in the bulk soil, *Ascomycota* was 66% followed by *Zygomycota* 28 % and Basidiomycota (3%)

### Host selection pressure, niche specification, and plant microbiome transmission

From source tracker analysis, a plant microbiome source prediction models were developed for both bacteria and fungi. For bacteria, bulk soil provided 87% of the rhizosphere microbial community. For root endophytes, rhizosphere and bulk soils served as a source of 52% and 30 % of the bacterial community while 18% came from an unknown source. Similarly in the leaf, root accounted for 53%, rhizosphere 14%, and bulk soil 10% of the bacterial community while 23% of the source could not be determined (Figure 1a).

**Figures 1.**
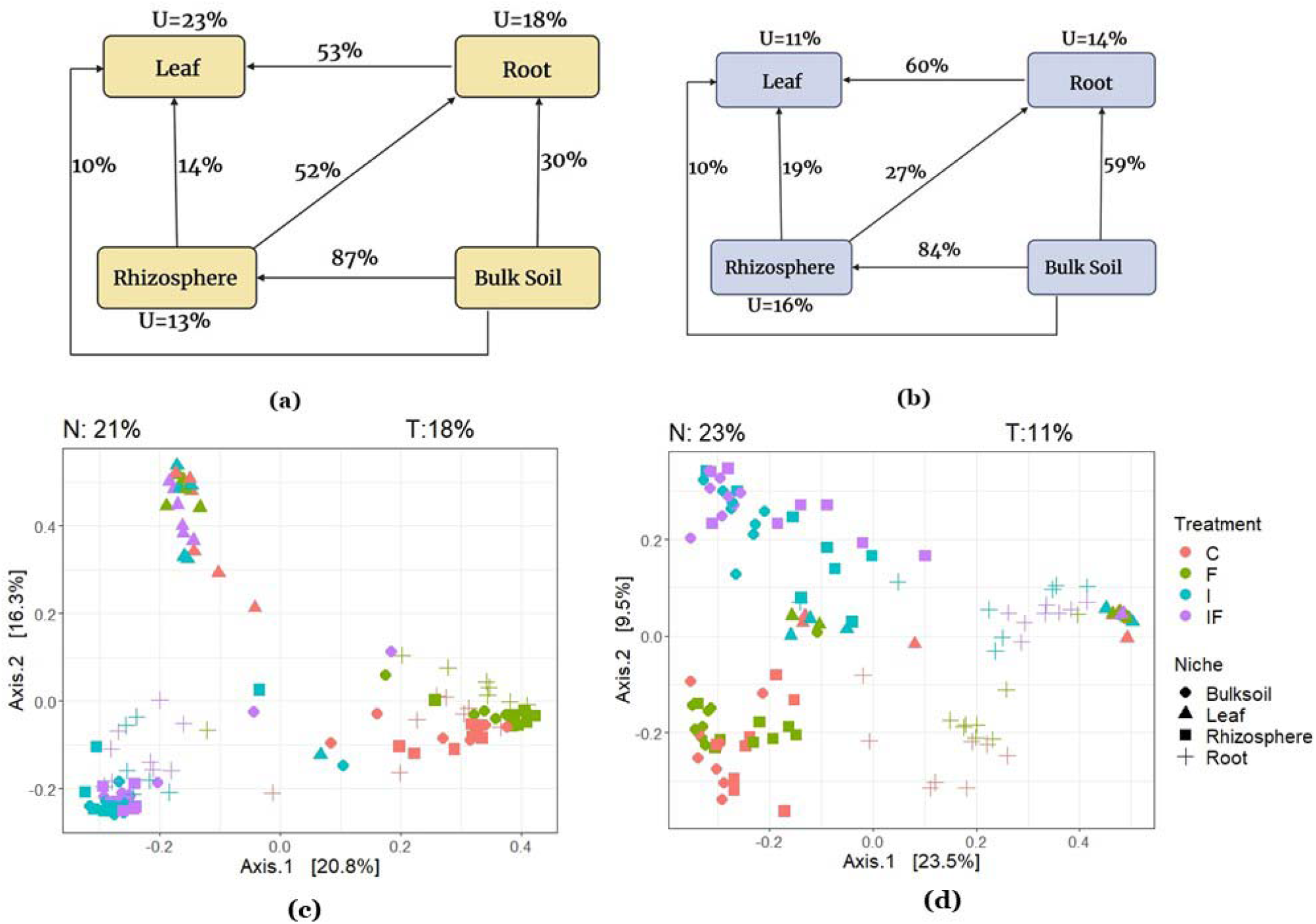

For fungi, similar trends of transmission were observed. Bulk soil served as a source for 84% of rhizosphere fungi. For the root microbiome 59% of members were from bulk soil and 27% from rhizosphere while 14% of fungi were unique to the root. In the case of leaf, root contributed 60%, rhizosphere 19%, and bulk soil provide 10% of the fungal community while the source for 11% of leaf mycobiome remain unaccounted (Figure 1b)

### Effect of land management practices on *E. saligna* microbiome

Irrigation and fertilization treatments had significant impacts on the alpha diversity of the bulk soil, rhizosphere, and root microbiome of bacteria. Fertilizer treatment reduced the diversity of all compartments except for leaves (Supplementary figure 7). For fungi, the alpha diversity of the rhizosphere and the bulk soil were significantly impacted by the fertilizer and irrigation treatment (Supplementary figure 8). Like bacteria, fertilizer decreased alpha diversity while irrigation increased it. Principal Co-ordinate and Permanova analyses showed that plant compartments had a greater effect on microbial community structure than management practices, for both bacteria (Figure 1c) and fungi (Figure 1d). The leaf microbiome formed a separate cluster for both bacteria and fungi and was not significantly affected by management practice, whereas the other niches responded significantly to land management practice. The effect of the niche was comparatively higher for fungal community (R^2^ =0.23834, P<0.001) compared to bacteria (R^2^ =0.20988, P<0.001) (Supplementary table 1). Management practices (fertilizers and irrigation) induced a clear separation in root, bulk soil and rhizosphere microbiome for both bacteria and fungi. However, the effect of management practice was stronger for bacteria R^2^=0.18294 (P <0.001) than fungi community (R^2^=0.1132; P <0.001)) (Supplementary table 1).. There was no significant effect of sampling time on the microbial community (R2=0.0093, P.value>0.5).

### Effect of land management practices on soil measurement and leaf traits

One-way ANOVA analysis revealed that management practices significantly influenced soil physiochemical properties including soil pH and moisture along with extractable nitrate (Table 2). Fertilizer treatment reduced pH (4.78125±0.23) while irrigation increased pH (6.51±0.31) as compared to control plots. Irrigation treatment also increased the soil moisture content (8.89 ± 0.83 %) as expected as compared to control plots

**Table 1:** Description of different niches used as sources for different sink niches.

| Serial No | Niche as Sink | Niche as Source |
| --- | --- | --- |
| 1 | Rhizosphere | Bulk Soil |
| 2 | Root | Rhizosphere and Bulk Soil |
| 3 | Leaf | Root, Rhizosphere, and Bulk Soil |

**Table 2:**
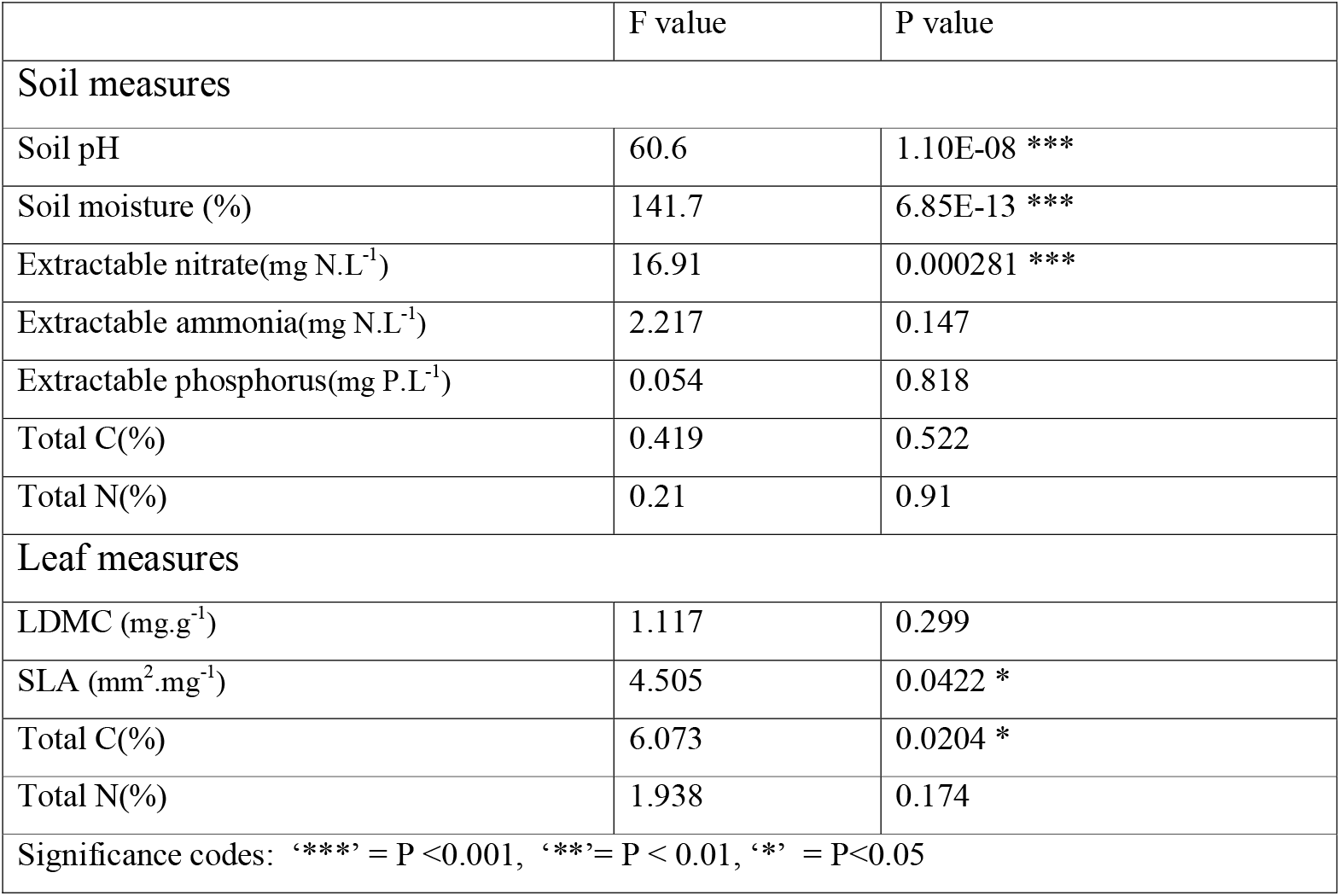
Table representing results from one-way ANOVA, soil pH, soil moisture, extractable nitrate, Specific Leaf Area (SLA), and leaf total C were significantly affected by land management practices. The last row of the table denotes the significance level (Significance codes: *** = P <0.001, **= P < 0.01, * = P<0.05)-.

|  | F value | P value |
| --- | --- | --- |
| **Soil measures** |  |  |
| Soil pH | 60.6 | 1.10E-08 *** |
| Soil moisture (%) | 141.7 | 6.85E-13 *** |
| Extractable nitrate(mg N.L<sup>-1</sup>) | 16.91 | 0.000281 *** |
| Extractable ammonia(mg N.L<sup>-1</sup>) | 2.217 | 0.147 |
| Extractable phosphorus(mg P.L<sup>-1</sup>) | 0.054 | 0.818 |
| Total C(%) | 0.419 | 0.522 |
| Total N(%) | 0.21 | 0.91 |
| **Leaf measures** |  |  |
| LDMC (mg.g<sup>-1</sup>) | 1.117 | 0.299 |
| SLA (mm<sup>2</sup>.mg<sup>-1</sup>) | 4.505 | 0.0422 * |
| Total C(%) | 6.073 | 0.0204 * |
| Total N(%) | 1.938 | 0.174 |
| Significance codes: '***' = $P < 0.001$, '**' = $P < 0.01$, '*' = $P < 0.05$ | | |

Among leaf traits, the Specific Leaf Area (SLA) and total C content of leaves were significantly affected by treatments. Soil total C and N contents were not significantly impacted by any of the management practices, while fertilizer and irrigation-fertilizer treatment increased extractable nitrate and ammonia along-with extractable phosphate content.

### Interactive impact of management practices, soil, and leaf properties, and microbial diversity

The generalized mixed model revealed a relationship between management practice, soil and leaf properties, and microbial diversity. When the effect of all factors were assessed together, the microbial diversity (Chao index) of all niches, responded differentially (Table 4). For bacteria, fertilizer treatment along with soil pH and soil nitrate concentrations significantly impacted the bacterial diversity of the bulk soil negatively. However, the rhizosphere bacterial community responded to soil pH, soil moisture, and irrigation treatment. Rhizosphere bacterial diversity negatively correlated with soil total N and soil ammonia. Soil total C along with irrigation and fertilizer treatment impacted the root bacterial diversity significantly. Root bacterial diversity positively correlated with soil moisture, irrigation-fertilizer and negatively with soil ammonia. Conversely, the bacterial diversity of leaves was neither significantly impacted by management practices nor by measured leaf traits.

**Table 3:** Mean values of various soil and leaf parameters studied across all four management practices. Letters in brackets denote the results obtained from Tukey’s posthoc test. ± indicates standard error.

| Soil factors | C | F | I | IF |
| --- | --- | --- | --- | --- |
| Soil pH | 5.02±0.10(a) | 4.78±0.23(a) | 6.63±0.31(b) | 6.56±0.14(b) |
| Soil Moisture (%) | 5.68±0.79(a) | 6.03±0.51(a) | 8.89±0.83(b) | 8.52±0.68(b) |
| Extractable Nitrate<br>(mg N.L<sup>-1</sup>) | 0.173±0.086(a) | 0.25±0.070(a) | 0.3100±.16(b) | 0.44±0.19(ab) |
| Extractable Ammonia<br>(mg N.L<sup>-1</sup>) | 0.21±0.065(ab) | 0.422±0.078(a) | 0.158±0.026(b) | 0.197±0.084(ab) |
| Extractable Phosphorus<br>(mg P.L<sup>-1</sup>) | 1.64±0.42(c) | 14.28±1.25(a) | 2.39±.029(c) | 6.266±0.96(b) |
| Total C(%) | 0.74±0.17(a) | 0.87±0.16(a) | 0.79±0.24(a) | 0.89±0.31(a) |
| Total N(%) | 0.07±0.011(a) | 0.16±0.06(a) | 0.08±0.03(a) | 0.14±0.03(a) |
| **Leaf measures** |  |  |  |  |
| LDMC(mg.g<sup>-1</sup>) | 247.51±25.92(a) | 261.00±39.42(a) | 252.28±25.01(a) | 273.75±26.40(a) |
| SLA(mm<sup>2</sup>.mg<sup>-1</sup>) | 12.84±1.98(a) | 13.36±2.02(a) | 13.52±1.88(a) | 14.84±1.51(a) |
| Total C% | 50.8±0.84(a) | 50.33±0.76(a) | 50.21±1.03(a) | 49.83±0.6(a) |
| Total N% | 2.45±0.50(a) | 2.57±0.51(a) | 2.61±0.56(a) | 2.92±0.42(a) |

**Table 4:** Results from generalised linear mixed model representing effect of management practices, soil measures and leaf traits over microbial alpha diversity (Significance codes: *** = P <0.001, **= P < 0.01, * = P<0.05)

| Niche | Soil<br>_pH | Soil_<br>Moistur<br>e | Soil<br>_N0<sub>3</sub> | Soil_<br>NH<sub>4</sub> | Soil_<br>PO<sub>4</sub> | Soil_<br>TotalN | Soil_<br>TotalC | I | F | IF | Leaf_<br>N | Leaf_<br>C | SLA | LDMC |
| --- | --- | --- | --- | --- | --- | --- | --- | --- | --- | --- | --- | --- | --- | --- |
| **Bacteria** |  |  |  |  |  |  |  |  |  |  |  |  |  |  |
| **Bulk Soil** | 0.02<br>0936<br>* | 0.139053 | 4.14<br>E-14<br>*** | 0.02068<br>4<br>* | 0.46456<br>5 | 0.31243<br>7 | 9.97E-11<br>*** | 0.01004<br>2<br>* | 0.2082<br>56 | 0.00034<br>2<br>*** | NA | NA | NA | NA |
| **Rhizosphere** | 0.30<br>602 | 0.010085<br>* | 0.00<br>023<br>7<br>*** | 0.00487<br>9<br>** | 0.03261<br>5<br>* | 0.01930<br>1<br>* | 0.135678 | 0.13537<br>5 | 0.0073<br>68<br>** | 0.31069<br>2 | NA | NA | NA | NA |
| **Root** | 0.01<br>98<br>* | 0.2114 | 0.68<br>49 | 0.0155<br>* | 0.5621 | 0.1146 | 0.3373 | 0.01<br>* | 0.3093 | 0.024<br>* | NA | NA | NA | NA |
| **Leaf** | NA | NA | Na | NA | NA | NA | NA | 0.84196 | 0.4854<br>9 | 0.23027 | 0.4818<br>4 | 0.4578 | 0.6298 | 0.5348 |
| **Fungi** |  |  |  |  |  |  |  |  |  |  |  |  |  |  |
| **Bulk Soil** | 0.03<br>2<br>* | 0.472 | 0.04<br>6<br>* | 0.339 | 0.124 | 0.751 | 0.562 | 0.431 | 0.0373<br>* | 0.602 | NA | NA | NA | NA |
| **Rhizosphere** | 0.03<br>63<br>* | 0.0404<br>* | 0.38<br>55 | 0.5579 | 0.6157 | 0.7351 | 0.171 | 0.0466<br>* | 0.5511 | 0.4009 | NA | NA | NA | NA |
| **Root** | 0.15<br>126 | 0.25365 | 0.68<br>088 | 0.05005<br>* | 0.21853 | 0.88805 | 0.00476<br>** | 0.00708<br>** | 0.4171<br>1 | 0.00781<br>** | NA | NA | NA | NA |
| **Leaf** | NA | NA | NA | NA | NA | NA | NA | 0.666 | 0.628 | 0.313 | 0.96 | 0.518 | 0.833 | 0.164 |

For fungi bulk soil diversity was correlated positively with extractable nitrate, and soil total C. Similarly, many soil-associated factors such as soil moisture, extractable nitrate, ammonia and phosphorus, and soil total had significant influence over fungal diversity. While root fungi were impacted by soil pH only. Unlike bacterial community, leaf fungal diversity was significantly positively correlated with SLA. The relation among microbial diversity, soil, and leaf-associated factors was further evaluated using Spearman correlations (Figure 2) which confirmed the above results.

**Figure 2.**
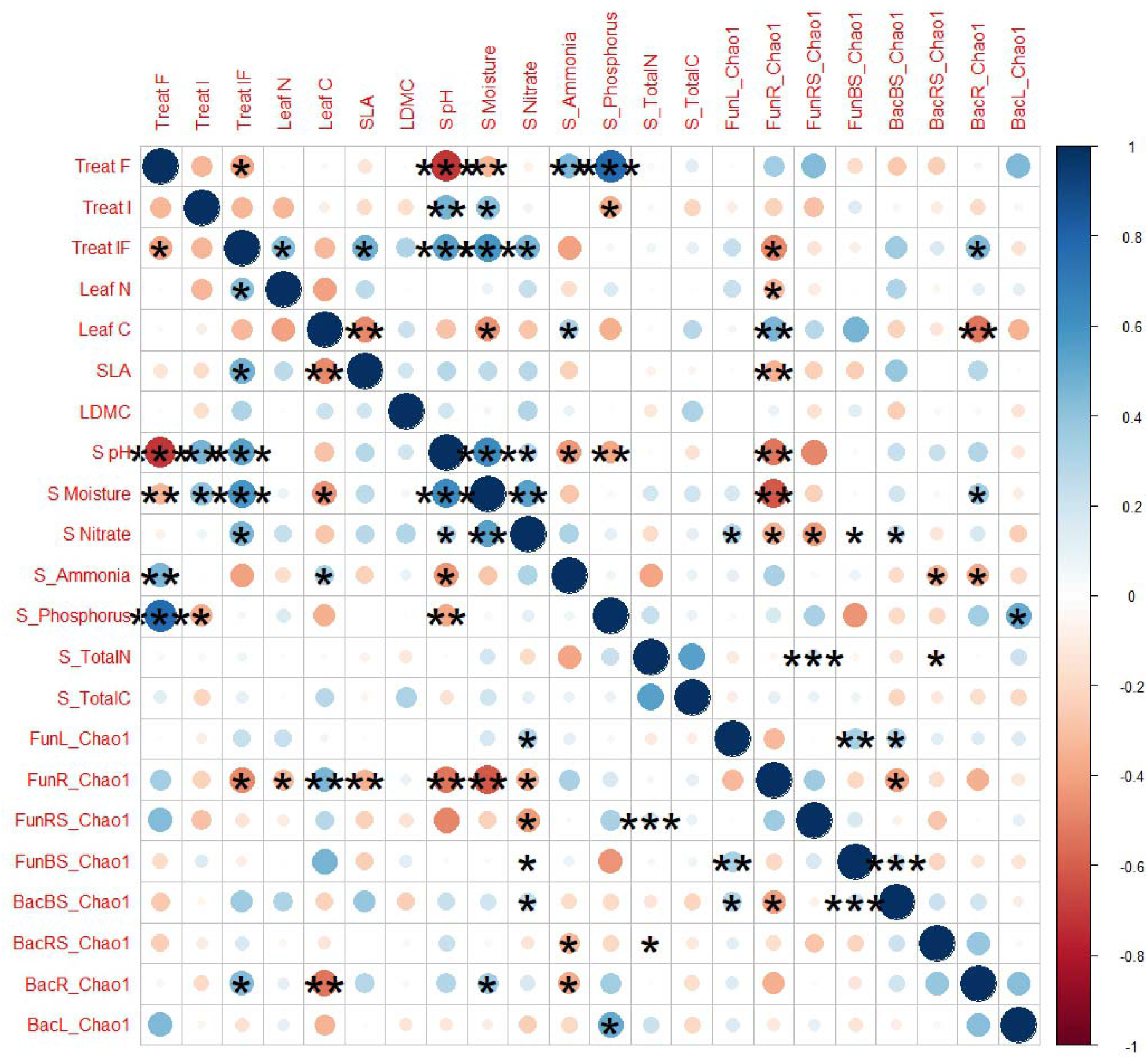

### Fungal OTUs are more responsive to land management practices than bacterial OTUs at the community level

Results from differential abundance analysis revealed different trends among bacteria and fungi across various niches(Figure 3 a and b). Fungal OTUs were more responsive toward land management practices. 16 differentially abundant bacterial OTUs (DABs) were identified in the leaf, 20 in the root, 25 in the rhizosphere, and 28 in bulk soil. Two DABs were common between and root, 1 between leaf-rhizosphere and leaf-bulk soil and 1 between leaf-rhizosphere and bulk soil. The number of shared DABs between bulk soil and rhizosphere was comparatively higher amounting to a total of 11, whereas root and rhizosphere shared 6 DABs among them (Figure 3 a).

**Figure 3.**
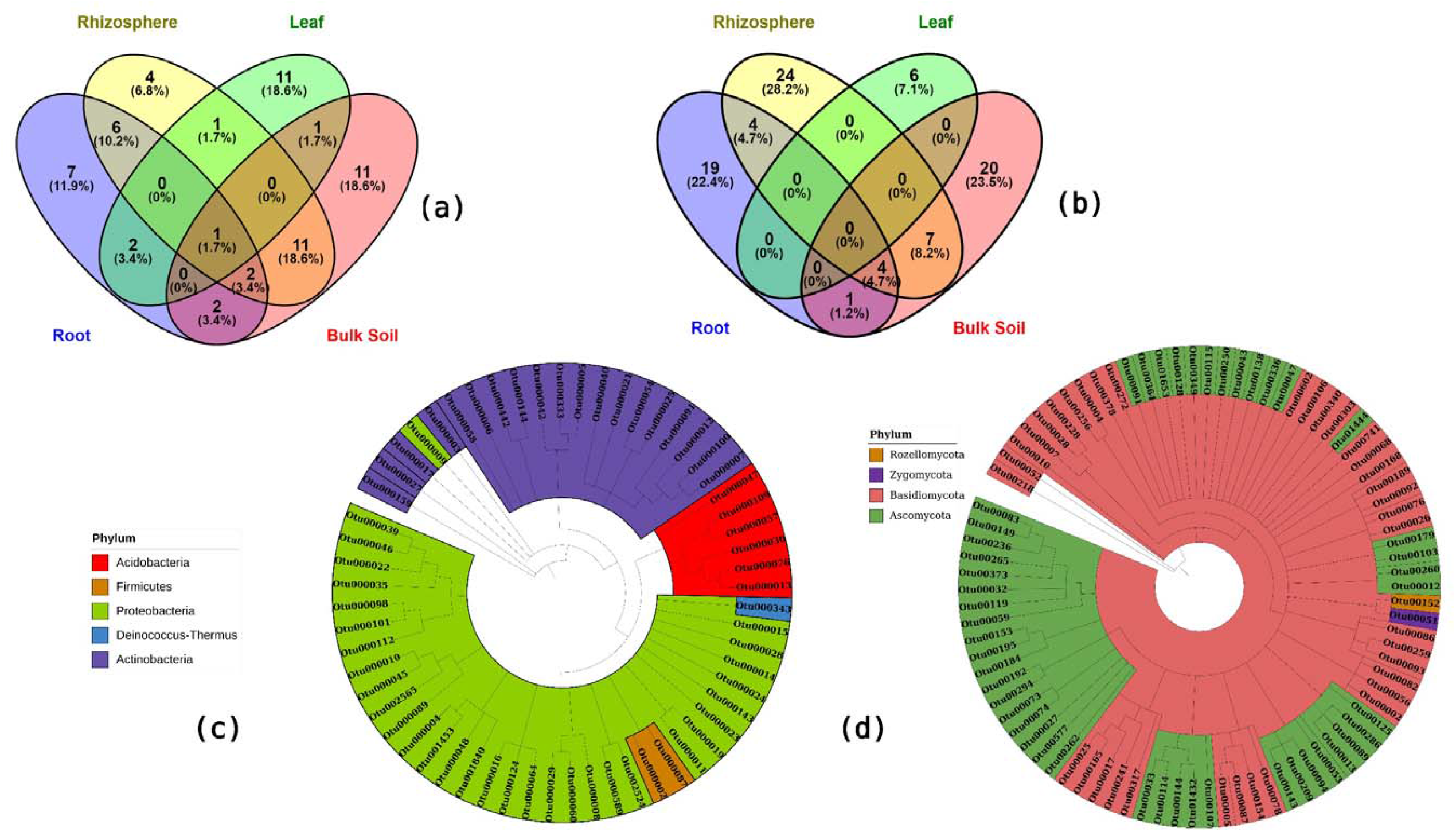

In leaves, bacterial OTUs belonging to *Alcaligenaceae* were comparatively more abundant in the fertilizer-treated plots, whereas OTUs from *Micromonoporineae* and *Bacillus* were more abundant in irrigation and irrigation-fertilizer-treated plots. (Supplementary figure 9). In the root, OTUS from the family *Acetobacteraceae* was less abundant in irrigated plots as compared to fertilized and control plots. Fertilized plots also had a lesser abundance of OTUs belonging to the family *Enterobacteriaceae*. While on the other hand, OTUs from *Xanthomonas, Streptomycinae*, and unclassified *Rhizobiales* were common in irrigation fertilizer combinations and irrigated plots for root, rhizosphere, and bulk soil (Supplementary figure 10,11,12). In addition, OTUs from the genus *Acidobacterium* were much more abundant in control and fertilized control plots across both rhizosphere and bulk soil.

Fertilized plots were characterized by an increased OTUs abundance from *Acetobacteraceae* and *Corneybacteraceae*, while irrigation led to an increased abundance of OTUs belonging to *Bacillus*

Fungi had a higher number of differentially abundant fungal OTUs (DAFs) as compared to bacteria. In the leaf a total of 6 DAFs were identified, for the root, there were a total of 28 DAFs, while at the same time, rhizosphere and bulk soil had 39 and 32 DAFs, respectively. Leaf had all exclusive DAFs, whereas 4 DAFs were shared between root-rhizosphere, root-bulk soil shared 1 DAFs, rhizosphere-bulk soil shared 7 DAFs and all three belowground compartments shared 4 DAFs among them. (Details of family level distribution of DAFs is in supplementary materials)

## Discussions

### Land management practices significantly affect below ground microbiome but not the leaf microbiome

Our findings suggest that while management practices strongly influenced bulk soil and rhizosphere microbial communities, plant endophytic microbiome, particularly leaf microbiome, were less impacted by management practices and soil properties. This suggests a strong role of host selection. Our finding is supported by previous reports of the fact that host-associated factors generally dominate leaf microbiome assembly compared to environmental factors (Wagner *et al*., 2016, Copeland and Paul Schulze-Lefert, 2020, Morella *et al*., 2020, Gao *et al*., 2020, Xiong *et al*., 2021). However, the soil serves as the main microbiome bank for the plants, and the source prediction model supported this for both root and leaf microbiome. The distinct insignificant response of leaf microbiome to management practices can be attributed to a unique environment and the hosts exert a stronger host selection mechanism (i.e. sequential filtration through rhizosphere, root and stems to reach to leaf) that potentially allow only specialized and plant recognized microbes to colonize the leaf (Compant *et al*., 2019; Trivedi *et al*., 2020). Leaf microbiome assembly is governed by host genetics, physiology and environment and their interactions. Host’s immune system and filtering mechanism allows only a certain category of microbes to colonize leaf, while soil acts as reservoir of competent microbes that can colonize plant tissues. The internal environment of the leaves provides a unique set of conditions for microbes and nutrient content, leaf chemistry and eco-physiological traits add up additional constraints for the microbe filtration (Compant et al., 2019; Trivedi et al., 2020). For example, leaf endophytes are less exposed to daily fluctuation in temperature and external inputs compared to soil/ rhizosphere microbiome. We observed that though management practices led to changes in total C of leaves and SLA, these changes did not alter diversity (alpha and beta) and composition of leaf microbiome. Additionally, we found that endophyte colonization in leaves was non-responsive to temporal effect. This suggests that plant endophytic microbiomes are subject to stronger host selection pressure which leads to a resilience towards regular fluctuation in the soil properties and hence the management effect is diluted as endophytic microbes moves from root to leaf (Xiong *et al*., 2020). In other words, soil properties effect soil microbiome more than root and leaf endophytes (Brown *et al*., 2020; Xiong *et al*., 2020). Our findings suggest that leaf endophytes are less susceptible to management practices whereas root endophytic microbes are still affected by land management practices, though the effect is much lower as compared to bulk and rhizosphere microbiome, supporting host sequential filtration theory.

The above discussion is supported by our source-tracker analysis that indicated that the bulk soil serves as a major source of endophyte microbes of *E. saligna*. This is supported by results from previous studies with crop species (Xiong *et al*., 2020, 2021). Our study provide evidence the sequential host filtering of microbiome in the following pattern-

Bulk Soil → Rhizosphere → Root → Stem → Leaf

Our results indicate that the host’s filtering mechanism becomes stronger as it moves from soil to leaf. This explains lower microbial diversity in root and leaf compared to bulk and rhizosphere soils. This finding is supported by previous studies which reported root has lower microbial diversity than the soil, while the leaf has much lower microbial diversity than the root (De Souza *et al*. 2016; Qian *et al*., 2019; Xiong *et al*., 2020; 2021). Further, our results suggest that while management practices have a strong impact on bulk soil microbial communities, plant endophytic microbiome, particularly leaf microbiome are less influenced by management practices, and soil properties suggesting a strong role of host selection. These findings are supported by findings of Wipf *et al*. (2021), where management practices affect soil microbiome more than the root microbiome of *Sorghum bicolor*. Similar findings were reported for drought affected rhizosphere more than the root microbiome (Fitzpatrick *et al*. 2018). Overall, our findings provide evidence that the host selection is main driver of endophyte assembly rather than management practices (Xiong *et al*. 2020), where leaf endophyte assembly was least affected by treatment as compared to bulk and rhizosphere soil microbiomes of wheat, maize, and barley crops.

On the other hand, a significantly high number of differentially abundant OTUs were observed for bacteria and fungi in response to management practices especially in below ground niches. The shared differentially abundant OTUs among different niches were quite low. For example, leaf fungi did not share any DAFs with other compartments, and similarly, shared bacterial DABs between the leaf and other compartments were also low, supporting the critical role of host selection in leaf colonization (Trivedi *et al* 2020; Xiong *et al*., 2021; Hamonts *et al* 2018). The dominance of genera such as *Bacillus, Pseudomonas, Stenotrophomonas, Pantoea, Propionibacterium*, and *Streptomyces* in the leaf supports the argument that host selection and microbes with specific abilities can colonize the leaf. Bacteria from some of these genera are known to colonize the leaves of various plant species, such as rice (Yang *et al*., 2020), maple trees (Wemheuer *et al*. 2019), wheat, and faba beans (Granzow *et al*., 2017). Our study advances this understanding by providing evidence from a long-term study that leaf microbiome assembly is comparatively more resistant to management practices as compared to soil and root microbiome.

The below-ground microbiome (i.e., bulk soil, rhizosphere, and root) responded significantly to land management practices. Soil-associated factors are known to serve as the major driving factor of soil microbiome assembly (Hossain and Sugiyama 2011; Xue *et al*., 2018) and rhizosphere microbiome likely due to direct contact with bulk soils (Xiong *et al*., 2020, 2021). Soil fertilization and irrigation impact soil edaphic properties, nutrient availability, and microbial diversity by enhancing microbial activities and host productivity which in turn leads to increased rhizodeposition and litter deposition. Though the rhizosphere effect plays an important role in shaping the rhizosphere microbiome (Compant *et al*. 2012, 2019: Li *et al*. 2022), these niches are more susceptible to change in the management practices than the above ground compartments, as they are exposed to treatments and associated physicochemical changes. Our results also support that in this environment (dryland) the microbiome assembly is driven by water availability than the nutrient enrichment through fertilizer treatment (Hu *et al*., 2015; Colombo et al. 2016).

### The application of fertilizer has a negative impact on the diversity of microbial communities in the soil and roots, while irrigation has the opposite effect

The bacterial community (both diversity and composition) responded more strongly to management practices than the fungal communities. The opposite was found for the effect of niche over microbiome assembly, as the effect of the niche was higher for fungal community compared to bacterial community. Fungal communities exhibited more resilience toward the management practices as they have complex genome and cellular structure as compared to bacteria. For example, fungi can acquire nutrients through mycelia which enables them to sustain in any temporary alteration in the environment whereas most bacteria have comparatively rapid life cycle which makes alteration in bacterial composition an easy task as compared to fungi. Similar observations were reported by De Vries *et al*. (2018) in which they found that bacterial networks were less stable than fungal network under drought condition. Similar findings were reported by Jiao *et al*. (2021) where soil bacterial communities and network were more affected than their fungal counterparts. Fungi have been reported to fluctuation in soil moisture and nutrient content as they can acquire nutrient through multiple sources (Hodge *et al*., 2001; Rousk *et al*., 2009).

Among the management treatments, the fertilizer treatment led to decreased soil and root bacterial diversity, whereas this trend was not evident for the fungal community. Conversely, irrigated plots had higher microbial diversity compared to the controlled plots. This trend supports previous studies from the same site (Colombo et al. 2016; Zheng *et al*., 2017) and from other regions (Hartmann *et al*., 2018; Frene *et al*., 2022). Interestingly, the clusters from PCoA analysis and ANOVA results on the Shannon diversity index suggested that across all below-ground microbiome, irrigation alone and irrigation-fertilizer treatments had similar profiles suggesting stronger impact of irrigation on microbial community compared to fertilization treatments. This can be explained by the facts that the fertilizer treatment, and irrigation have opposite effects on the soil edaphic properties as well soil nutrients. Increased soil pH and soil moisture due to irrigation has provided a comparatively better niche for microbes, hence the diversity was comparatively higher in these plots. Water availability has been reported to have a positive impact on biological processes and microbial functions in arid region and particularly at this site by previous studies support the results (Hu *et al*, 2015: Martins *et al*., 2015; Colombo et al. 2016). The relative abundance of copiotrophs belonging to families such as *Bacillaceae, Hyphomicrobiaceae, Caulobacteraceae* increased in the irrigated plots supporting that this ecosystem is limited by water and less so for nutrient availability.

Fertilizer treatment interferes with soil microbiome assembly in multiple ways Fertilizers typically contain high levels of nitrogen, phosphorus, and potassium that can lead to an increase in the growth of certain bacteria and fungi that proliferate in high nutrients environments(Alori et al., 2017, while suppressing the growth of other microbes that are sensitive to high nutrient conditions. In addition to the decreased pH, fertilizer treatment led to low soil moisture content and comparatively very high amount of extractable ammonia and phosphate. This explains why there was reduced microbial diversity given that it promoted the growth of acidophiles. For example, in bacteria, fertilizer treatment was associated with an increased abundance of oligotrophs belonging to families such as *Acidobacterium, Acetobacteraceae, Catenulisporineae*, and *Edaphobacter* in both soil (bulk and rhizosphere) and root microbiome families. The abundance of these families has been reported to negatively correlate with soil pH (Lauber *et al*., 2009; Rousk *et al*., 2010; Hegyi *et al*., 2021).

In the fungal community, the fertilizer treatment led to an enrichment of saprotrophs such as *Helicoma sp, Mortierell sp* and *Trechispora sp*, along with Ecto-mycorrhizal fungi like *Cenococcum sp*. This supports the finding of Zheng *et al*. (2017) in which they reported that management practices enriched saprotrophic and mycorrhizal fungi. Especially *Mortrierella sp*. has been reported to thrive in fertilizer-treated low-pH soil as it can produce acid during metabolic processes (Han *et al*., 2010, Wen *et al*., 2020).

Overall, our study demonstrates that leaf endophyte diversity is largely resilient to management practices, such as irrigation and fertilization, with only minor changes in community composition observed. In contrast, the diversity and community structure of below-ground microbiome were significantly impacted by management practices. These results suggest that leaf microbiome is primarily shaped by host selection and are protected from external fluctuations in physico-chemical properties, while bulk and rhizosphere soils are strongly responsive to management practices. Therefore, future studies on microbial response to environmental factors should explicitly consider compartment niches, as microbiome of different compartments can have a differential impact on host health and functions.

## Acknowledgement

This research was funded by Australian Research Council. PS received PhD scholarship from Western Sydney University. EE acknowledges DECRA fellowship from Australian Research councils. The site was established with support from the Australian Greenhouse Office grant 0506/0085 and subsequently by the Commonwealth Department of Climate Change, with additional funding from the NSW Department of Environment and Climate Change (grant T07/CAG/16). Authors acknowledge Ramesha Jayaramaiah and Burhan Amiji for their help in sampling.

