## Supplementary table 1, Supplementary table 2,Supplementary table 3 for "Host selection has stronger impact on leaf microbiome assembly compared to land-management practices"

**Supplementary files:**

**Supplementary figures:**

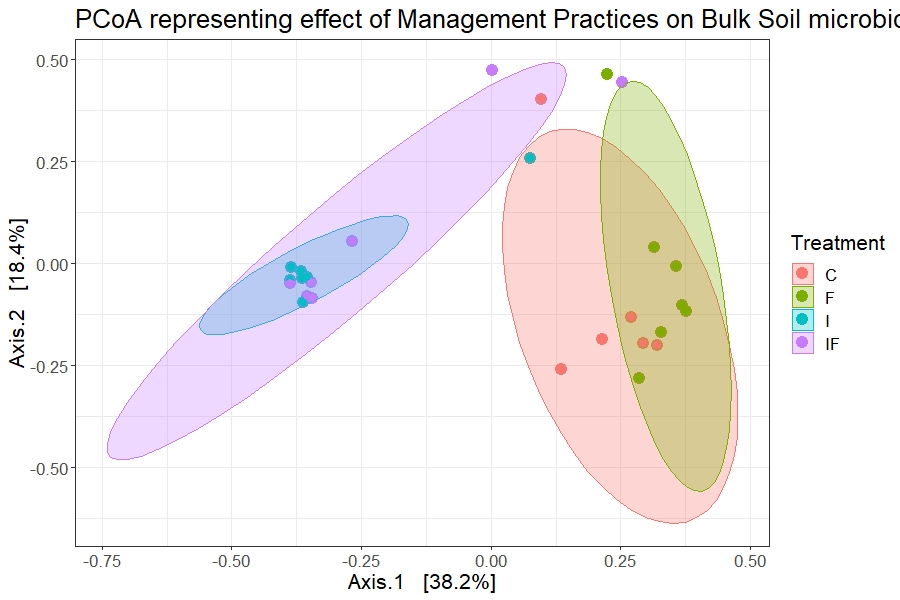

**Supplementary figure 1**: PCoA plot representing the effect of management practices on bulk soil bacterial microbiome when rarified at 10000 reads (Control: C, Fertilizer: F, Irrigation: I and, Irrigation & Fertilizer: IF)

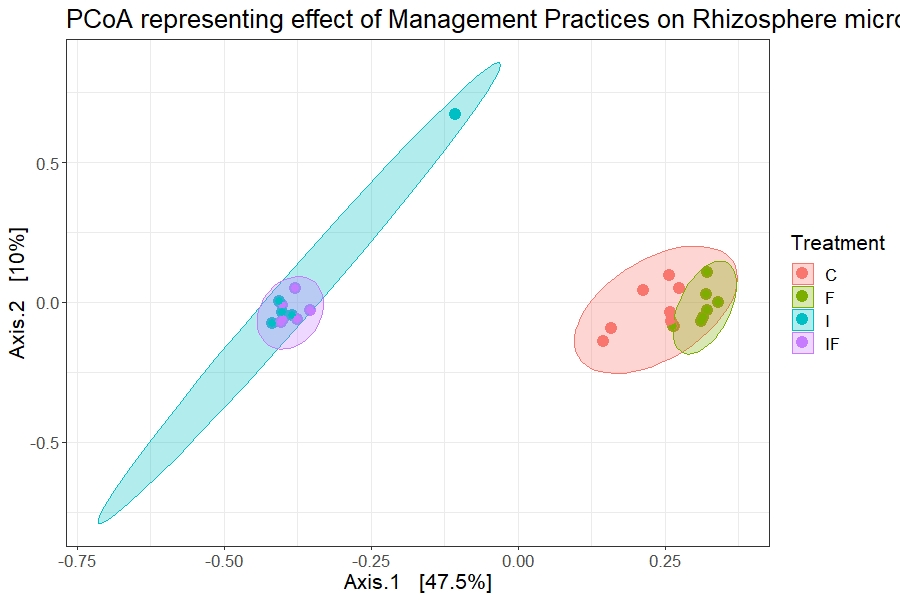

**Supplementary figure 2**: PCoA plot representing the effect of management practices on rhizosphere soil bacterial microbiome when rarified at 10000 reads (Control: C, Fertilizer: F, Irrigation: I and, Irrigation & Fertilizer: IF)

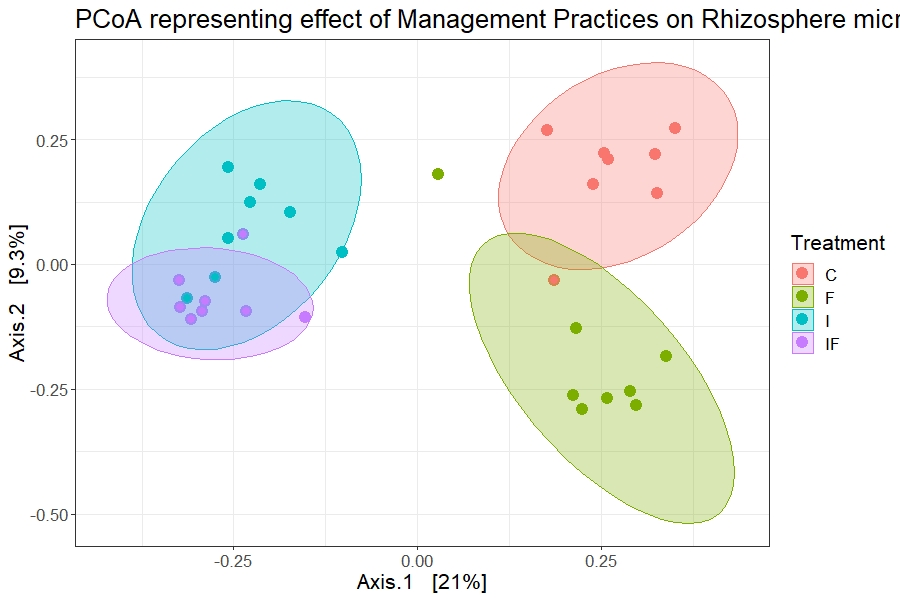

**Supplementary figure 3**: PCoA plot representing the effect of management practices on bulk soil fungal microbiome when rarified at 10000 reads (Control: C, Fertilizer: F, Irrigation: I and, Irrigation & Fertilizer: IF)

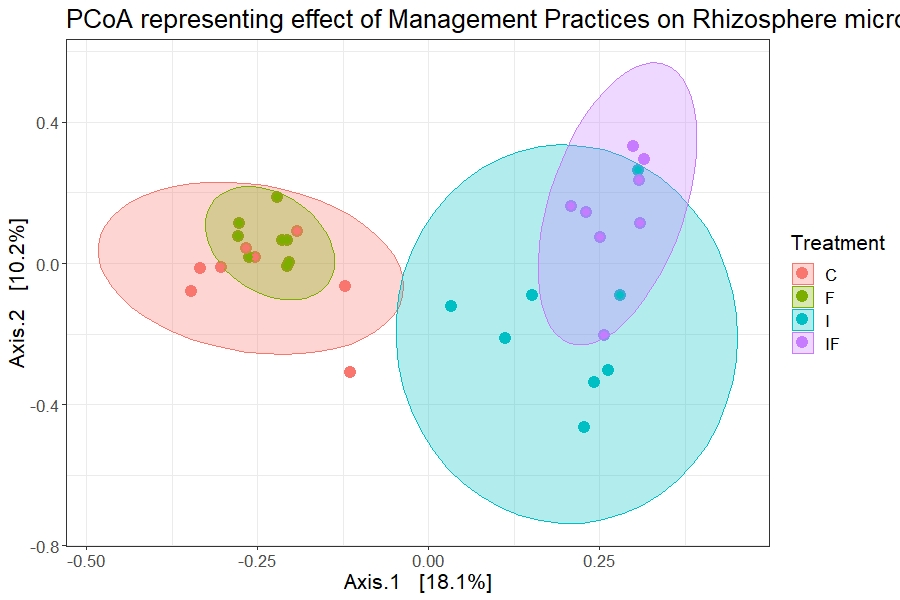

**Supplementary figure 4**: PCoA plot representing the effect of management practices on rhizosphere soil fungal microbiome when rarified at 10000 reads (Control: C, Fertilizer: F, Irrigation: I and, Irrigation & Fertilizer: IF

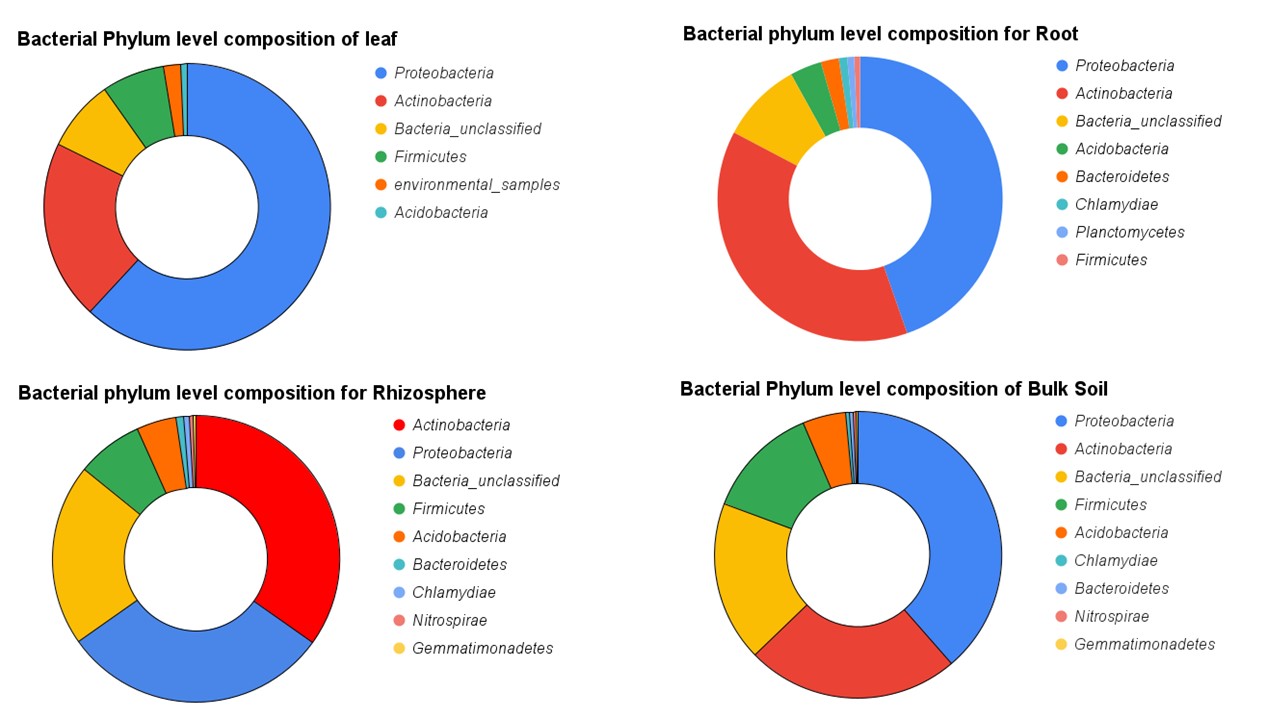

**Supplementary figure 5**: Pie charts representing bacterial phylum level composition for all four niches, i.e., leaf, root, rhizosphere, and bulk soil

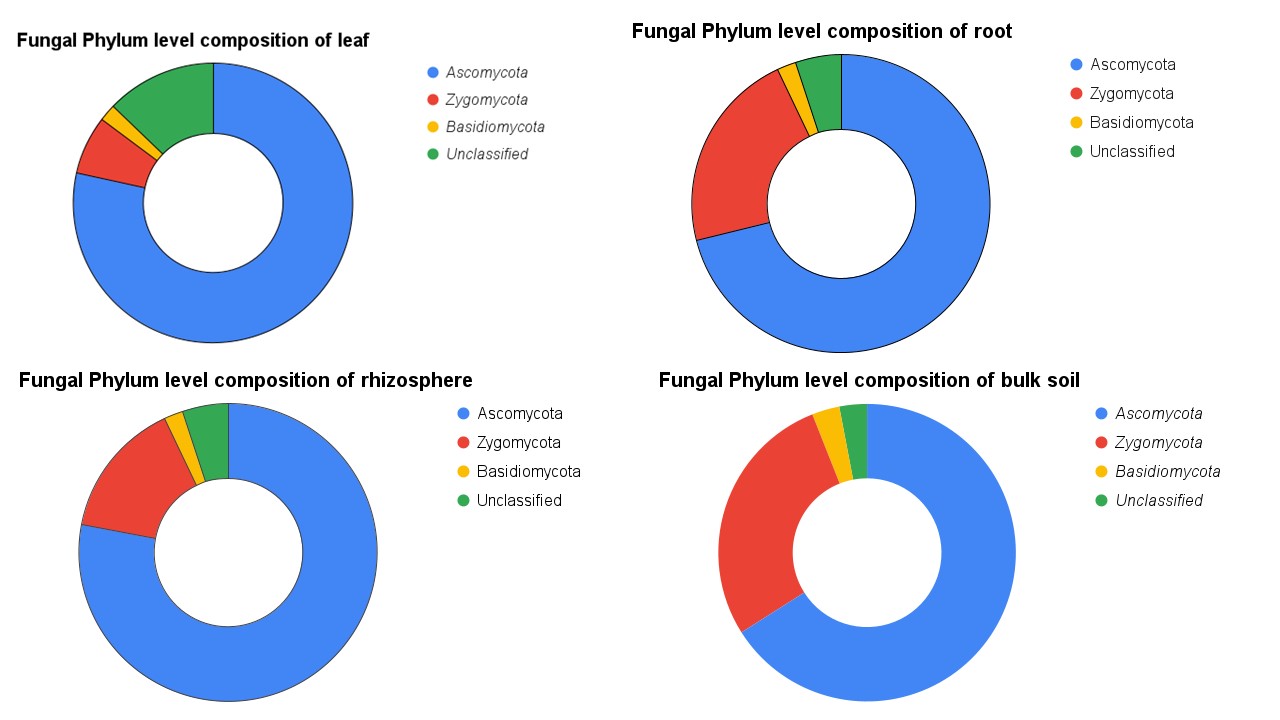
 **Supplementary figure 6**: Pie charts representing fungal phylum level composition for all four niches, i.e., leaf, root, rhizosphere, and bulk soil

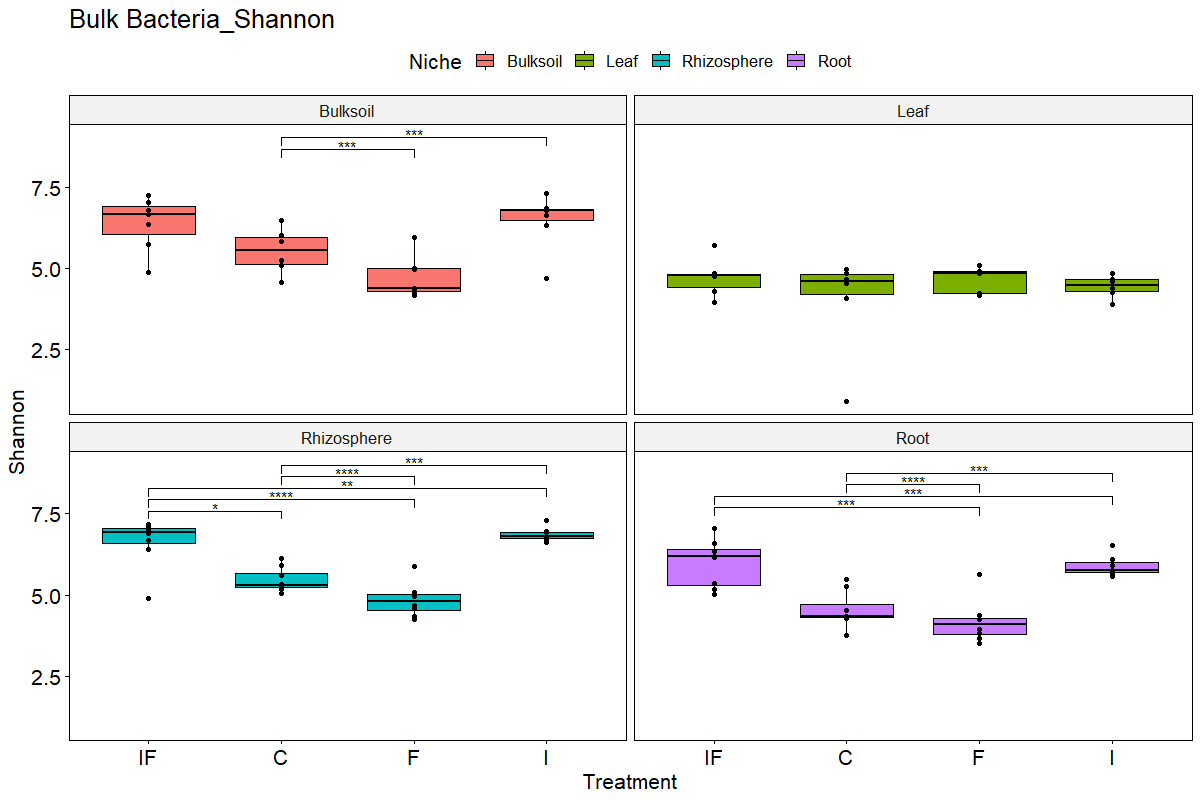

**Supplementary figure 7:** Boxplots representing bacterial Shannon diversity indexes and impact of land management practices across different niches (Control: C, Fertilizer: F, Irrigation: I, and, Irrigation & Fertilizer: IF) Significance codes: P.Value< 0 ‘***’ 0.001 ‘**’ 0.01 ‘*’ 0.05.

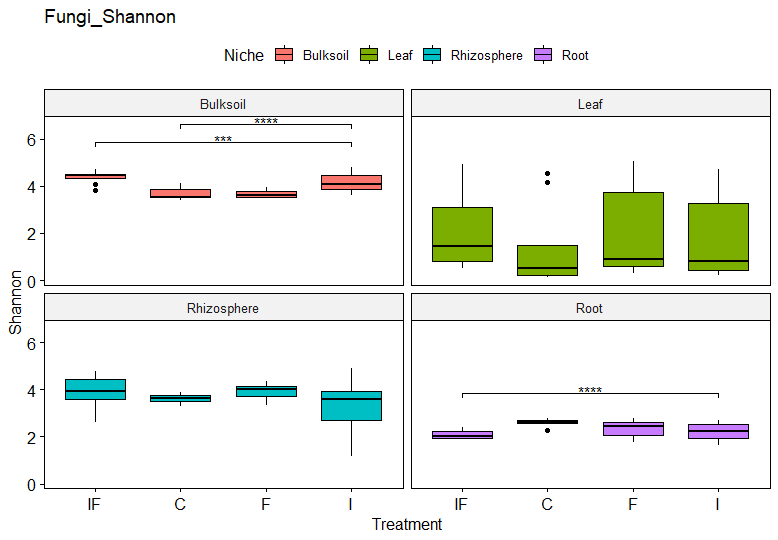

**Supplementary figure 8**: Boxplots representing fungal Shannon diversity indexes and impact of land management practices across different niches (Control: C, Fertilizer: F, Irrigation: I, and, Irrigation & Fertilizer: IF) Significance codes: P.Value< 0 ‘***’ 0.001 ‘**’ 0.01 ‘*’ 0.05.

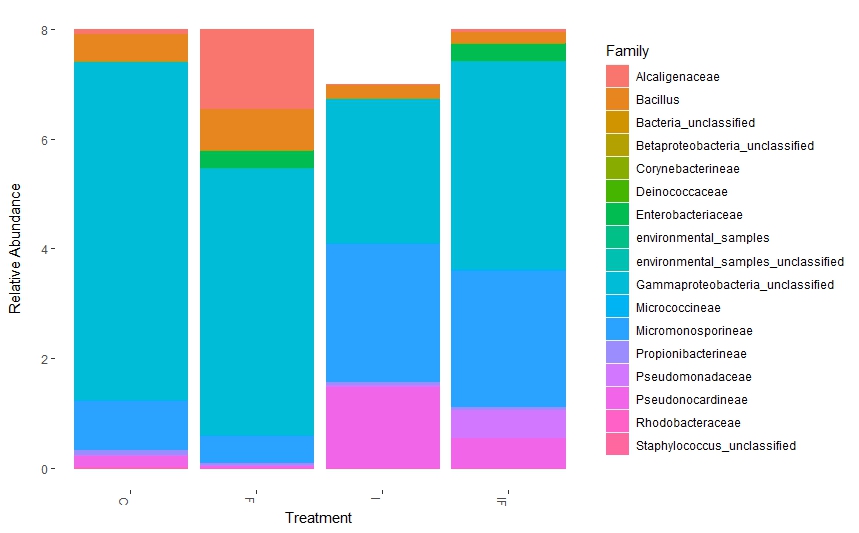

**Supplementary figure 9**: Bar plot representing the family level composition of differentially abundant bacterial OTUs for leaf across all treatments i.e., management practices (Control: C, Fertilizer: F, Irrigation: I and, Irrigation & Fertilizer: IF)

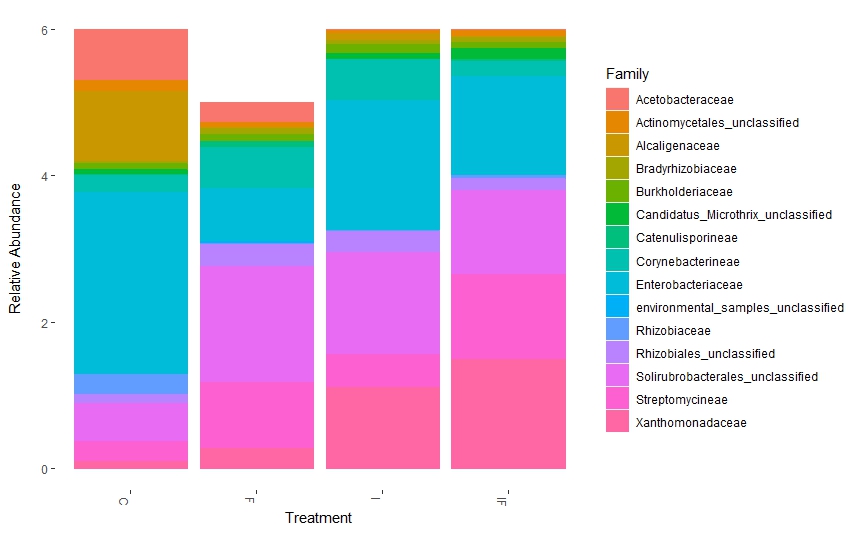

**Supplementary figure 10:** Bar plot representing family level composition of differentially abundant bacterial OTUs for root across all treatments i.e., management practices (Control: C, Fertilizer: F, Irrigation: I and, Irrigation & Fertilizer: IF)

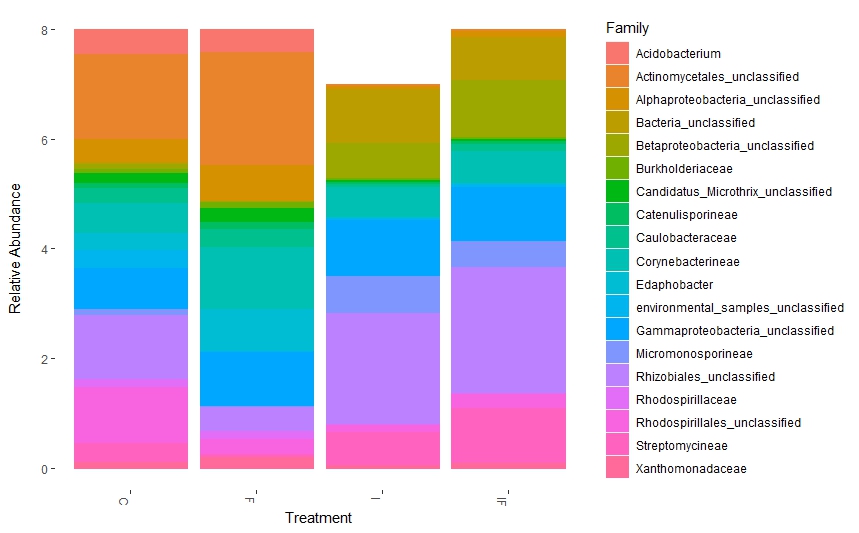

**Supplementary figure 11**: Bar plot representing the family level composition of differentially abundant bacterial OTUs for rhizosphere across all treatments i.e., management practices (Control: C, Fertilizer: F, Irrigation: I and, Irrigation & Fertilizer: IF)

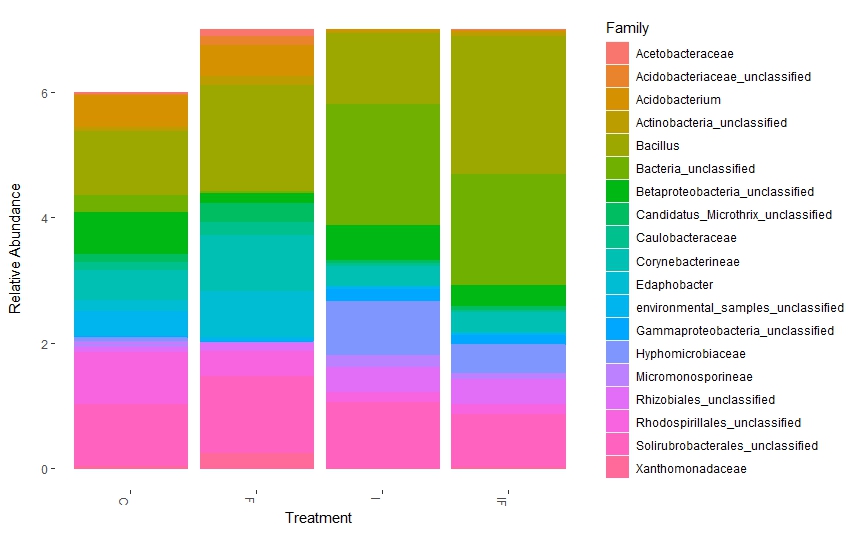

**Supplementary figure 12**: Bar plot representing the family level composition of differentially abundant bacterial OTUs for bulk soil across all treatments i.e., management practices (Control: C, Fertilizer: F, Irrigation: I and, Irrigation & Fertilizer: IF)

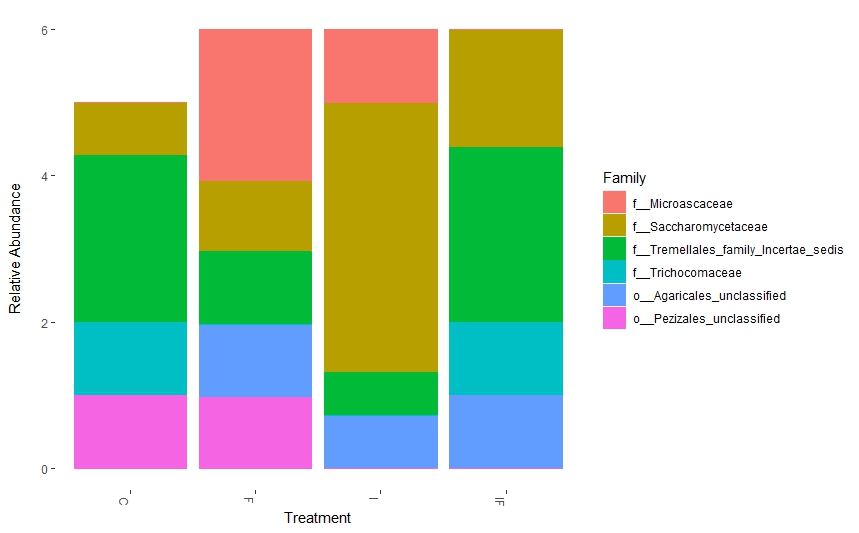

**Supplementary figure 13**: Bar plot representing the family level composition of differentially abundant fungal OTUs for leaf across all treatments i.e., management practices (Control: C, Fertilizer: F, Irrigation: I and, Irrigation & Fertilizer: IF)

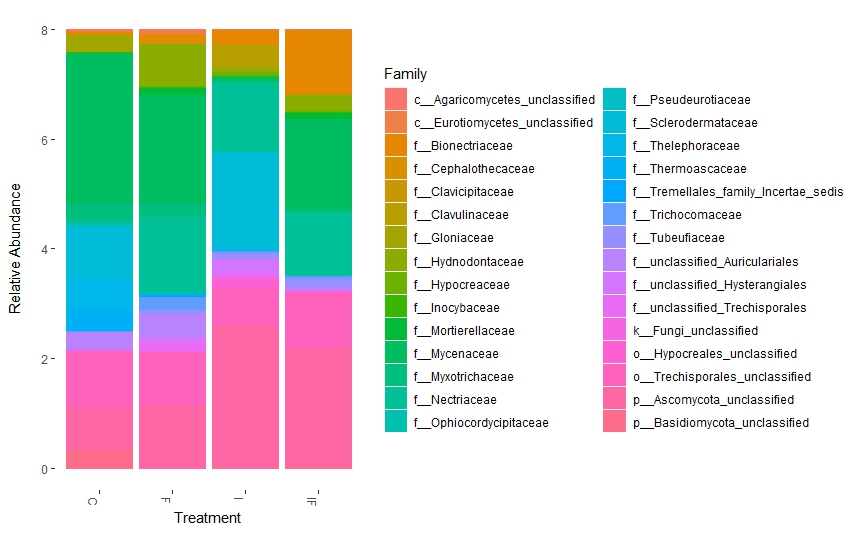

**Supplementary figure 14**: Bar plot representing the family level composition of differentially abundant fungal OTUs for root across all treatments i.e., management practices (Control: C, Fertilizer: F, Irrigation: I and, Irrigation & Fertilizer: IF)

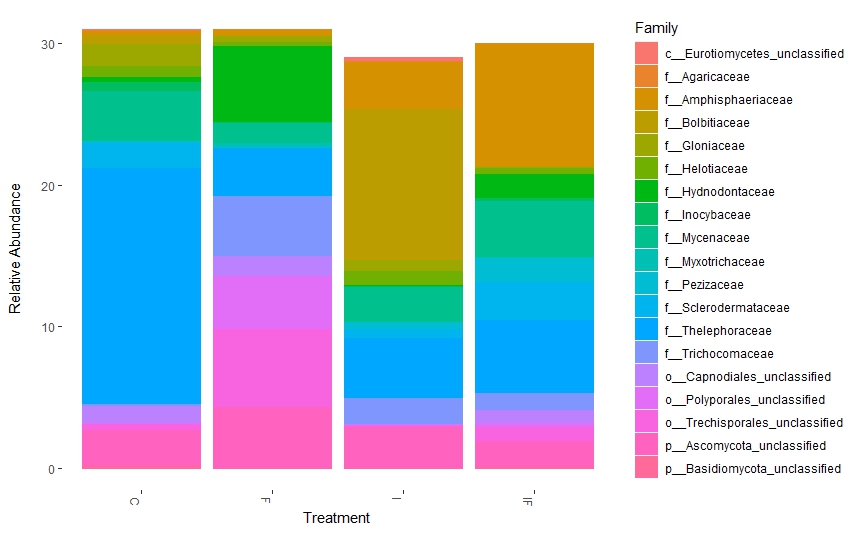

**Supplementary figure 15**: Bar plot representing the family level composition of differentially abundant fungal OTUs for rhizosphere across all treatments i.e., management practices (Control: C, Fertilizer: F, Irrigation: I and, Irrigation & Fertilizer: IF)

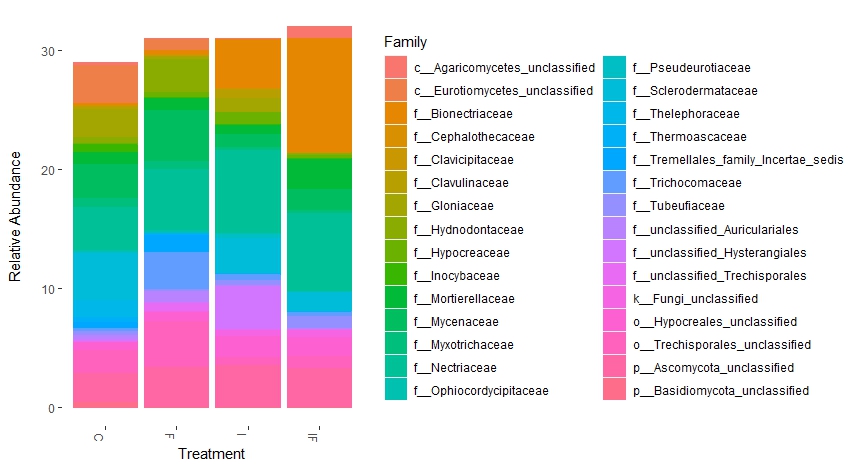

**Supplementary figure 16**: Bar plot representing the family level composition of differentially abundant fungal OTUs for bulk soil across all treatments i.e., management practices (Control: C, Fertilizer: F, Irrigation: I and, Irrigation & Fertilizer: IF)

**Supplementary table**

**Supplementary table 1:**

PERMANOVA table representing the effect of management practices over different niches for bacteria and fungi. (Significance codes: P.Value< 0 *** 0.001 ** 0.01 * 0.05.)

|  | F.Model | R^2^ | Pr(>F) |
| --- | --- | --- | --- |
| Bacteria | | | |
| Leaf | 1.0686 | 0.14437 | 0.314 |
| Root | 7.2195 | 0.44511 | 1e-04 *** |
| Rhizosphere | 14.319 | 0.61405 | 0.001 *** |
| Bulk Soil | 7.1825 | 0.4837 | 0.001 *** |
| Fungi | | | |
| Leaf | 0.45638 | 0.04662 | 0.966 |
| Root | 3.52 | 0.27386 | 1e-04 *** |
| Rhizosphere | 4.3298 | 0.3169 | 1e-04 *** |
| Bulk Soil | 4.8639 | 0.3426 | 1e-04 *** |

**Supplementary Table 2** : Taxonomy and niche based details for Differentially Abundant Bacteria (DAFs)

| OTUID | *Phylum* | *Genus* | Niche |
| --- | --- | --- | --- |
| Otu000002 | *Firmicutes* | *Bacillus benzoevoran* | Leaf and Bulk Soil |
| Otu000003 | *Actinobacteria* | *Solirubrobacterales_unclassified* | Leaf and Rhizosphere |
| Otu000004 | *Actinobacteria* | *Mycobacteriaceae* | Leaf, Root and Rhizosphere |
| Otu000005 | *Proteobacteria* | *Betaproteobacteria_unclassified* | Leaf and Root |
| Otu000006 | *Acidobacteria* | *Edaphobacter_unclassified* | Leaf and Root |
| Otu000007 | *Proteobacteria* | *Pedomicrobium* | Leaf |
| Otu000008 | *Proteobacteria* | *Xanthomonadaceae_unclassified* | Leaf |
| Otu000010 | *Bacteria_unclassified* | *Bacteria_unclassified* | Leaf |
| Otu000011 | *Proteobacteria* | *Rhizobiales_unclassified* | Leaf |
| Otu000012 | *Actinobacteria* | *Micromonosporaceae* | Leaf |
| Otu000013 | *Proteobacteria* | *Rhodospirillales_unclassified* | Leaf |
| Otu000014 | *Proteobacteria* | *Caulobacteraceae_unclassified* | Leaf |
| Otu000015 | *Actinobacteria* | *Candidatus_Microthrix_unclassified* | Leaf |
| Otu000016 | *Bacteria_unclassified* | *Bacteria_unclassified* | Leaf |
| Otu000017 | *Bacteria_unclassified* | *Bacteria_unclassified* | Leaf |
| Otu000019 | *Acidobacteria* | *environmental_samples* | Leaf |
| Otu000021 | *Proteobacteria* | *Betaproteobacteria_unclassified* | Root |
| Otu000022 | *Bacteria_unclassified* | *Bacteria_unclassified* | Root |
| Otu000023 | *Proteobacteria* | *Rhodospirillales_unclassified* | Root |
| Otu000024 | *Acidobacteria* | *environmental_samples_unclassified* | Root |
| Otu000025 | *Actinobacteria* | *Actinobacteria_unclassified* | Root |
| Otu000027 | *Acidobacteria* | *environmental_samples* | Root |
| Otu000028 | *Proteobacteria* | *Gammaproteobacteria_unclassified* | Root |
| Otu000029 | *Proteobacteria* | *Xanthomonadaceae_unclassified* | Root and Rhizosphere |
| Otu000030 | *Acidobacteria* | *Acidobacteriaceae_unclassified* | Root and Rhizosphere |
| Otu000033 | *Proteobacteria* | *Acetobacteraceae_unclassified* | Root and Rhizosphere |
| Otu000035 | *Acidobacteria* | *environmental_samples* | Root and Rhizosphere |
| Otu000039 | *Bacteria_unclassified* | *Bacteria_unclassified* | Root and Rhizosphere |
| Otu000040 | *Firmicutes* | *environmental_samples* | Root and Rhizosphere |
| Otu000045 | *Proteobacteria* | *Pantoea* | Root and Bulk Soil |
| Otu000046 | *Actinobacteria* | *Pseudonocardiaceae* | Root and Bulk Soil |
| Otu000047 | *Proteobacteria* | *Pseudomonas* | Root, Rhizosphere and Bulk Soil |
| Otu000048 | *Actinobacteria* | *Propionibacteriaceae* | Root, Rhizosphere and Bulk Soil |
| Otu000054 | *Firmicutes* | *Staphylococcus_unclassified* | Rhizosphere and Bulk Soil |
| Otu000057 | *Proteobacteria* | *Alcaligenaceae_unclassified* | Rhizosphere and Bulk Soil |
| Otu000058 | *Actinobacteria* | *Micrococcaceae* | Rhizosphere and Bulk Soil |
| Otu000060 | *Deinococcus-Thermus* | *Deinococcus* | Rhizosphere and Bulk Soil |
| Otu000064 | *Actinobacteria* | *Intrasporangiaceae* | Rhizosphere and Bulk Soil |
| Otu000076 | *Bacteria_unclassified* | *Bacteria_unclassified* | Rhizosphere and Bulk Soil |
| Otu000087 | *Proteobacteria* | *Gammaproteobacteria_unclassified* | Rhizosphere and Bulk Soil |
| Otu000089 | *Proteobacteria* | *Gammaproteobacteria_unclassified* | Rhizosphere and Bulk Soil |
| Otu000091 | *environmental_samples* | *environmental_samples_unclassified* | Rhizosphere and Bulk Soil |
| Otu000098 | *Proteobacteria* | *Betaproteobacteria_unclassified* | Rhizosphere and Bulk Soil |
| Otu000100 | *Actinobacteria* | *Streptomycetaceae* | Rhizosphere and Bulk Soil |
| Otu000101 | *Actinobacteria* | *Actinomycetales_unclassified* | Rhizosphere |
| Otu000109 | *Proteobacteria* | *Gammaproteobacteria_unclassified* | Rhizosphere |
| Otu000112 | *Proteobacteria* | *Rhizobiales_unclassified* | Rhizosphere |
| Otu000124 | *Actinobacteria* | *Actinomycetales_unclassified* | Rhizosphere |
| Otu000143 | *Proteobacteria* | *Alphaproteobacteria_unclassified* | Bulk Soil |
| Otu000144 | *Proteobacteria* | *Gammaproteobacteria_unclassified* | Bulk Soil |
| Otu000159 | *Actinobacteria* | *Actinospicaceae* | Bulk Soil |
| Otu000333 | *Proteobacteria* | *Burkholderiaceae_unclassified* | Bulk Soil |
| Otu000343 | *Proteobacteria* | *Rhodospirillaceae_unclassified* | Bulk Soil |
| Otu000442 | *Actinobacteria* | *Actinomycetales_unclassified* | Bulk Soil |
| Otu000589 | *Proteobacteria* | *Bradyrhizobiaceae_unclassified* | Bulk Soil |
| Otu001453 | *Actinobacteria* | *Actinomycetales_unclassified* | Bulk Soil |
| Otu001840 | *Proteobacteria* | *Acetobacteraceae_unclassified* | Bulk Soil |
| Otu002524 | *Proteobacteria* | *Acidocella* | Bulk Soil |
| Otu002565 | *Proteobacteria* | *Stenotrophomonas* | Bulk Soil |

**Supplementary table 3**: Taxonomy and niche based details for Differentially Abundant Fungi (DAFs)

| OTUID | *Phylum* | *Genus* | Niche |
| --- | --- | --- | --- |
| Otu00002 | *Basidiomycota* | *Thelephora* | root and rhizosphere |
| Otu00004 | *Basidiomycota* | *unclassified_Basidiomycota* | root and rhizosphere |
| Otu00005 | *Basidiomycota* | *Scleroderma* | root and rhizosphere |
| Otu00007 | *Basidiomycota* | *unclassified_Trechisporales* | Rhizosphere |
| Otu00010 | *Basidiomycota* | *Trechisporales_unclassified* | Bulk Soil |
| Otu00012 | *Ascomycota* | *Geomyces* | Rhizosphere |
| Otu00013 | *Ascomycota* | *Oidiodendron* | Rhizosphere |
| Otu00017 | *Basidiomycota* | *Cruentomycena* | Bulk Soil |
| Otu00025 | *Basidiomycota* | *Mycena* | Rhizosphere |
| Otu00026 | *Basidiomycota* | *Hydnangiaceae_unclassified* | Root |
| Otu00027 | *Ascomycota* | *Fusarium* | Bulk Soil |
| Otu00028 | *Basidiomycota* | *Trechispora* | Bulk Soil, Rhizosphere and Root |
| Otu00032 | *Ascomycota* | *Bionectriaceae_unclassified* | Bulk Soil |
| Otu00033 | *Ascomycota* | *Trichocomaceae_unclassified* | Bulk Soil, Rhizosphere and Root |
| Otu00043 | *Ascomycota* | *Cenococcum* | Bulk Soil, Rhizosphere and Root |
| Otu00047 | *Ascomycota* | *Phaeohelotium* | Rhizosphere |
| Otu00051 | *Zygomycota* | *Mortierella* | Bulk Soil and Rhizosphere |
| Otu00052 | *Basidiomycota* | *Trechisporales_unclassified* | Bulk Soil |
| Otu00053 | *Ascomycota* | *Ascomycota_unclassified* | Bulk Soil |
| Otu00056 | *Basidiomycota* | *Thelephoraceae_unclassified* | Root |
| Otu00059 | *Ascomycota* | *Ascomycota_unclassified* | Bulk Soil and Root |
| Otu00068 | *Basidiomycota* | *Polyporales_unclassified* | Root |
| Otu00073 | *Ascomycota* | *Codinaeopsis* | rootand rhizosphere |
| Otu00074 | *Ascomycota* | *Ascomycota_unclassified* | Root |
| Otu00076 | *Basidiomycota* | *Descomyces* | Rhizosphere |
| Otu00078 | *Basidiomycota* | *Pisolithus* | Bulk Soil, Rhizosphere and Root |
| Otu00082 | *Basidiomycota* | *Thelephoraceae_unclassified* | Bulk Soil and Rhizosphere |
| Otu00083 | *Ascomycota* | *Hypocreaceae_unclassified* | Root |
| Otu00086 | *Basidiomycota* | *unclassified_Auriculariales* | Bulk Soil |
| Otu00087 | *Basidiomycota* | *Scleroderma* | Bulk Soil |
| Otu00089 | *Ascomycota* | *Oidiodendron* | Bulk Soil |
| Otu00091 | *Ascomycota* | *Saccharomyces* | Leaf |
| Otu00092 | *Basidiomycota* | *Agaricomycetes_unclassified* | Root |
| Otu00093 | *Basidiomycota* | *Auriculariales_family_Incertae_sedis_unclassified* | Root |
| Otu00094 | *Ascomycota* | *Capnodium* | Rhizosphere |
| Otu00103 | *Ascomycota* | *Ascomycota_unclassified* | Bulk Soil and Rhizosphere |
| Otu00106 | *Basidiomycota* | *Cryptococcus* | Bulk Soil and Rhizosphere |
| Otu00107 | *Ascomycota* | *Ascomycota_unclassified* | Bulk Soil |
| Otu00114 | *Ascomycota* | *Talaromyces* | Rhizosphere |
| Otu00115 | *Ascomycota* | *Helicoma* | Bulk Soil |
| Otu00119 | *Ascomycota* | *Hypocreales_unclassified* | Bulk Soil |
| Otu00125 | *Ascomycota* | *Fusidium* | Root |
| Otu00128 | *Ascomycota* | *Phaeomoniella* | Root |
| Otu00138 | *Ascomycota* | *Ascomycota_unclassified* | Rhizosphere |
| Otu00143 | *Ascomycota* | *Devriesia* | Rhizosphere |
| Otu00144 | *Ascomycota* | *Trichocomaceae_unclassified* | Bulk Soil and Rhizosphere |
| Otu00149 | *Ascomycota* | *Hypocrea* | Bulk Soil |
| Otu00153 | *Ascomycota* | *Lasiosphaeriaceae_unclassified* | Root |
| Otu00154 | *Basidiomycota* | *Scleroderma* | Bulk Soil and Rhizosphere |
| Otu00165 | *Basidiomycota* | *Mycena* | Root |
| Otu00168 | *Basidiomycota* | *Inocybaceae_unclassified* | Bulk Soil and Rhizosphere |
| Otu00179 | *Ascomycota* | *Phoma* | Rhizosphere |
| Otu00184 | *Ascomycota* | *Ascomycota_family_Incertae_sedis_unclassified* | Rhizosphere |
| Otu00189 | *Basidiomycota* | *Clavulina* | Bulk Soil |
| Otu00192 | *Ascomycota* | *Immersidiscosia* | Rhizosphere |
| Otu00195 | *Ascomycota* | *Sordariomycetes_unclassified* | Root |
| Otu00209 | *Ascomycota* | *Capnodiales_unclassified* | Rhizosphere |
| Otu00218 | *Basidiomycota* | *Trechisporales_unclassified* | Root |
| Otu00228 | *Basidiomycota* | *Trechisporales_unclassified* | Rhizosphere |
| Otu00236 | *Ascomycota* | *Ophiocordyceps* | Rhizosphere |
| Otu00241 | *Basidiomycota* | *Mycenaceae_unclassified* | Root |
| Otu00250 | *Ascomycota* | *Dothideales_unclassified* | Rhizosphere |
| Otu00256 | *Basidiomycota* | *unclassified_Trechisporales* | Bulk Soil |
| Otu00152 | *Rozellomycota* | *unclassified_Rozellomycota* | Root |
| Otu00259 | *Basidiomycota* | *Auricularia* | Root |
| Otu00260 | *Ascomycota* | *Pseudeurotium* | Bulk Soil |
| Otu00262 | *Ascomycota* | *unclassified_Thermoascaceae* | Bulk Soil |
| Otu00265 | *Ascomycota* | *Ophiocordycipitaceae_unclassified* | Bulk Soil |
| Otu00272 | *Basidiomycota* | *Agaricaceae_unclassified* | Rhizosphere |
| Otu00286 | *Ascomycota* | *Oidiodendron* | Rhizosphere |
| Otu00294 | *Ascomycota* | *Amphisphaeriaceae_unclassified* | Rhizosphere |
| Otu00303 | *Basidiomycota* | *Agaricomycetes_unclassified* | Root |
| Otu00317 | *Basidiomycota* | *Mycenaceae_unclassified* | Root |
| Otu00336 | *Ascomycota* | *Ascomycota_unclassified* | Rhizosphere |
| Otu00340 | *Basidiomycota* | *Agaricomycetes_unclassified* | Root |
| Otu00349 | *Ascomycota* | *Pseudocercospora* | Rhizosphere |
| Otu00364 | *Ascomycota* | *Pezizaceae_unclassified* | Rhizosphere |
| Otu00373 | *Ascomycota* | *Clavicipitaceae_unclassified* | Bulk Soil |
| Otu00378 | *Basidiomycota* | *Leucoagaricus* | Rhizosphere |
| Otu00577 | *Ascomycota* | *Cephalotheca* | Bulk Soil |
| Otu00602 | *Basidiomycota* | *Filobasidiella* | Leaf |
| Otu00741 | *Basidiomycota* | *Agaricales_unclassified* | Leaf |
| Otu01432 | *Ascomycota* | *Aspergillus* | Leaf |
| Otu01442 | *Ascomycota* | *unclassified_Microascaceae* | Leaf |
| Otu01653 | *Ascomycota* | *Pezizales_unclassified* | Leaf |

**Family level distribution of differentially abundant fungal OTUs;**

At the family level, the leaf mycobiome of the fertilized plots had a higher abundance of *Microscaceae* and Unclassified *Pezizales* while fertilizer treatment resulted in an increased abundance of *Saccharomycetaceae* (Supplementary figure 13). For root, Fertilizer treatment led to increased colonization by *Inocybaceae*, while irrigation treatment led to an increased abundance of *Thelephoraceae* (Supplementary figure 14). In the case of rhizosphere fertilizer treatment led to an increased abundance of *Hydnodontaceae* and *Trichocomaceae*. Contrastingly, irrigation treatment was characterized by an increased abundance of *Agaricaceae* and *Amphisphaeriaceae* (Supplementary 15). For bulk soil, *Inocybaceae* were abundant in fertilized plots, whereas irrigated plots were characterized by an increased abundance of *Bionectriaceae*(Supplementary figure 16).
